# Revisiting object contextual cueing: A replication study on implicit learning

**DOI:** 10.1101/2025.07.30.667636

**Authors:** Zsófia Anna Gaál, Petia Kojouharova, István Czigler, Boglárka Nagy

## Abstract

Object contextual cueing is an implicit learning paradigm in which the repeated co-occurrence of certain objects facilitates target detection, even when their spatial arrangement varies. In two experiments – where participants performed a visual search task in which the target was accompanied by either consistently repeated or randomly varying distractors – we attempted to replicate Chun and Jiang’s (1999) study. In Experiment 1 (N = 25, M = 20.8 years), although participants became faster and more accurate over time, no significant difference emerged between repeated and new configurations. Thus, using the same experimental design, we were unable to replicate the object contextual cueing effect. In Experiment 2, we tested whether search strategy influences the effect by instructing one group to adopt an active search strategy and another to use a passive one (N = 20 per group, M = 21.8 and 21.7 years, respectively; active: actively search for the target, passive: let the unique item pop out). However, the manipulation had no effect: the contextual cueing effect did not emerge in either group. Importantly, within each group we observed substantial individual differences – some participants showed a facilitation effect, others no effect, and some even showed a reverse pattern. These findings suggest that object contextual cueing may not be a robust or universal phenomenon and may instead depend on individual cognitive styles or learning tendencies.

**Highlights:**

- Replication of Chun and Jiang’s (1999) object cueing study was unsuccessful.
- Active vs. passive strategy instructions had no effect on object contextual cueing.
- Large individual differences emerged in the contextual cueing effect.

## 1.. INTRODUCTION

In our visual environment, we constantly search for something – whether it is the TV remote, a phone charger, or a friend in a theatre lobby. In these situations, various stimuli in the environment and the regularities among them guide our attention and help us to find what or whom we are looking for. If certain objects or people frequently appear together, after just a few encounters, we become faster at locating our target.

Consequently, context plays a crucial role in guiding our attention. When considering objects, context refers to their arrangement and dynamics when perceived as the target in view. It encompasses the multitude of stimuli that constantly surround us, from which we have to extract relevant information for our actions. In young adults, context facilitates object recognition (Mudrik et al., 2014) and localization, especially when detecting a stimulus is difficult, such as in blurred vision or ambiguous conditions (Oliva & Torralba, 2007).

This phenomenon greatly depends on implicit learning, the process of acquiring knowledge without awareness or intent, which plays a significant role in shaping perception and cognitive processes. During implicit learning, we unconsciously extract recurring patterns from the stimuli in our environment (Goujon et al., 2015). This form of learning has several advantages: it is highly robust, resistant to interference (Zellin et al., 2014), less affected by negative psychological states (Rathus et al., 1994) or high cognitive load (Vickery et al., 2010), and once established, it can persist over long periods (Chun & Jiang, 2003). Due to its parallel and non-sequential processing, implicit learning allows for the processing of large amounts of information, enabling the detection of complex patterns (Seger, 1994).

Compared to explicit learning, where new information is consciously acquired, implicit learning relies on distinct neural mechanisms (Reber, 2013). These mechanisms have the advantage of being less susceptible to aging effects (Robin & Moscovitch, 2017) and often remain intact in various psychiatric and neurological disorders (Eldridge et al., 2002; Danion et al., 2001). This raises the possibility that implicit learning could be utilized as compensation for individuals whose working memory limitations hinder explicit learning, such as in older adults.

Implicit learning also has its drawbacks. It is less flexible, not easily generalizable to new situations, and the acquired knowledge is difficult to verbalize, limiting its conscious application (Goujon et al., 2015). However, some situations do not require explicit awareness, and implicit processes can even support explicit learning. For instance, older adults may sometimes forget why they walked into the kitchen. However, if certain objects are consistently present – such as a radio, calendar, puzzle book, and pen on the table – these items could draw their attention to an always-present pill organizer, helping them realize they came to take their medication.

Our study focuses on implicit learning, specifically on how context can guide attention. One well-studied form of this is spatial contextual cueing (Chun & Jiang, 1998, Kojouharova et al., 2023). In this type of task, participants have to locate a target stimulus (a “T” in the original experiment) among distractors (various configurations of “L” letters). Some spatial configurations repeat over multiple trials, while others appear only once. Research has shown that participants quickly learn the relationship between repeated spatial configurations and the target’s location, allowing them to direct their attention more efficiently and respond faster.

However, Chun and Jiang (1999) conducted another experiment where the spatial arrangement of the stimuli was not fixed. Instead, in repeated configurations, only the objects themselves (symmetrical shapes) remained consistent with the target stimulus. Even in this case, participants were able to locate the target faster in repeated configurations compared to configurations presented only a single time – a phenomenon known as object cueing. Since objects frequently change locations in daily life, we likely rely more on a mechanism that recognizes co-occurring objects rather than fixed spatial positions, allowing for quicker identification.

Surprisingly, despite its potential significance, object cueing effect has not been systematically replicated, nor have the factors influencing its effects been explored. Our study aimed to replicate Chun and Jiang’s (1999) original object cueing paradigm and extend it by using not only meaningless geometric shapes but also faces, which are highly relevant stimuli in everyday life.

## 2. METHODS

### 2.1. Participants

University students participated in two experiments, and all received course credit or financial compensation for their participation. They had normal or corrected-to-normal vision (at least 5/5 on a version of the Snellen charts), and they reported no neurological or psychiatric disorders. The study was approved by local ethical review committee. Written informed consent was obtained from all participants included in the study in accordance with the Declaration of Helsinki.

### 2.2. Stimuli and procedure

Each participant sat in a comfortable chair within a soundproof and electrically shielded room. We used Cogent 2000 (Cogent 2000 team, 2003) and Cogent Graphics (Romaya, 2008) toolboxes in MATLAB (MathWorks, Inc., 2015, Natick, MA, USA) for stimulus presentation and response recording. The stimuli were presented on a 24-inch monitor (BenQ, resolution 1920 × 1080, 60 Hz refresh rate) from 1.2 m distance, and responses were made by using a mouse (Logitech M150 wireless mouse, Lausanne, Switzerland). Participants first completed an object cueing task based on Chun and Jiang (1999), followed by an old/new recognition test. The details are provided in the experiment sections below.

### 2.3. Data analysis

For the statistical analyses, we used JASP 0.19.3 (JASP Team, 2025). We analysed the behavioural data of the blocks (details provided in the experiment sections), including mean reaction time (RT) and mean error rate (misses and incorrect responses) to examine how contextual information was acquired at the behavioural level as the task progressed. Repeated measures ANOVAs were conducted with *Block* (1-6) and *Configuration* (REPEATED/NEW) as within-subject factors. In post hoc analysis of Experiment 1, and in Experiment 2, *Group* (CC/ NO CC and ACTIVE/PASSIVE, respectively) was included as a between-subject factor. Greenhouse-Geisser correction was applied, when necessary, with p-values reflecting these corrected values. Effect sizes were calculated as partial eta square (η^2^_p_), and post hoc analysis was conducted using Tukey’s honestly significant difference (HSD) test.

RT analysis included only correct responses and excluded trials with RT faster than 150 ms. The mean error rate included trials where the target was not detected within the specified time window and trials with incorrect target localization. We defined the emergence of the contextual cueing effect occurring when the average RT difference in Blocks 2-6 between the NEW and REPEATED conditions exceeded zero.

To test whether participants recognized the repetition of the presented configurations during the contextual cueing task, we conducted an old/new recognition test. We assessed the accuracy of responses by measuring the rates of correct (hits and correct rejections) and erroneous (misses and false alarms) responses for both REPEATED and NEW configurations. In this context, hits and misses correspond to yes/no responses to REPEATED configurations, while false alarms and correct rejections correspond to yes/no responses to NEW configurations. We used the Two-High Threshold Model by Snodgrass and Corvin (1988) to adjust for guessing.

Using these measures, we calculated two behavioural indices. The first is the discrimination index (Pr), which indicates the probability that a configuration (REPEATED or NEW) exceeds the old/new recognition threshold. It is calculated as Pr = HR - FAR, where HR represents the hit rate and FAR the false alarm rate. A Pr value of 0 suggests decisions were made by chance; therefore, we compared the Pr values to 0 using single-sample t-tests to determine if repeated configurations led to lasting representations in our participants.

The second index is the bias index (Br), which reflects the likelihood of a yes (old) response under uncertainty. It is calculated as Br = FAR/[1 - (HR - FAR)]. A Br value between 0.5 and 1 indicates a liberal bias, where more yes/old responses are given under uncertainty, while a Br value between 0 and 0.5 indicates a conservative bias, where more no/new responses are given. To identify the bias type in our sample, we compared the mean Br values to 0.5 using single-sample t-tests.

## 4. EXPERIMENT 1

### 4.1. METHODS

Twenty-five younger adults participated in the study (19 women, M = 20.8 years, SD = 2.6, all right-handed – the original Chun and Jiang study consisted of 17 students).

In this replication of Chun and Jiang’s (1999) experiment, the target was identified as the only object with vertical axis symmetry, while distractor stimuli had symmetry along axes deviating from the vertical. This method effectively designated a search target without specifying its exact shape. The experimental design is shown in Figure 1. In total, 96 objects were generated by MATLAB, 16 of which were targets, and 80 were distractors. Sixteen different stimuli served as distractors in each of five orientation groups; they were symmetric around the 30°, 60°, 90°, 120°, and 150° axes. All stimuli (1.52° x 1.14°) were presented in white on a grey background. In each trial, 1 target and 10 distractor stimuli (comprising 2 different shapes from all 5 of the distractor orientation groups) were randomly placed within an 8 × 6 grid (40° x 30°). We had REPEATED and NEW conditions, but unlike in spatial contextual cueing studies, where target and distractor locations were fixed, both the target’s position and the distractor arrangement were randomized in every trial. In the REPEATED (consistent-mapping) condition, each target was consistently paired with a specific distractor set – meaning the same shapes were presented together, although their position was not fixed – and this pairing remained the same throughout the experiment. In the NEW (variable-mapping) condition, the pairings between targets and distractors were randomized in each block, meaning that always different shaped distractors were presented with the given target.

**Figure 1.**
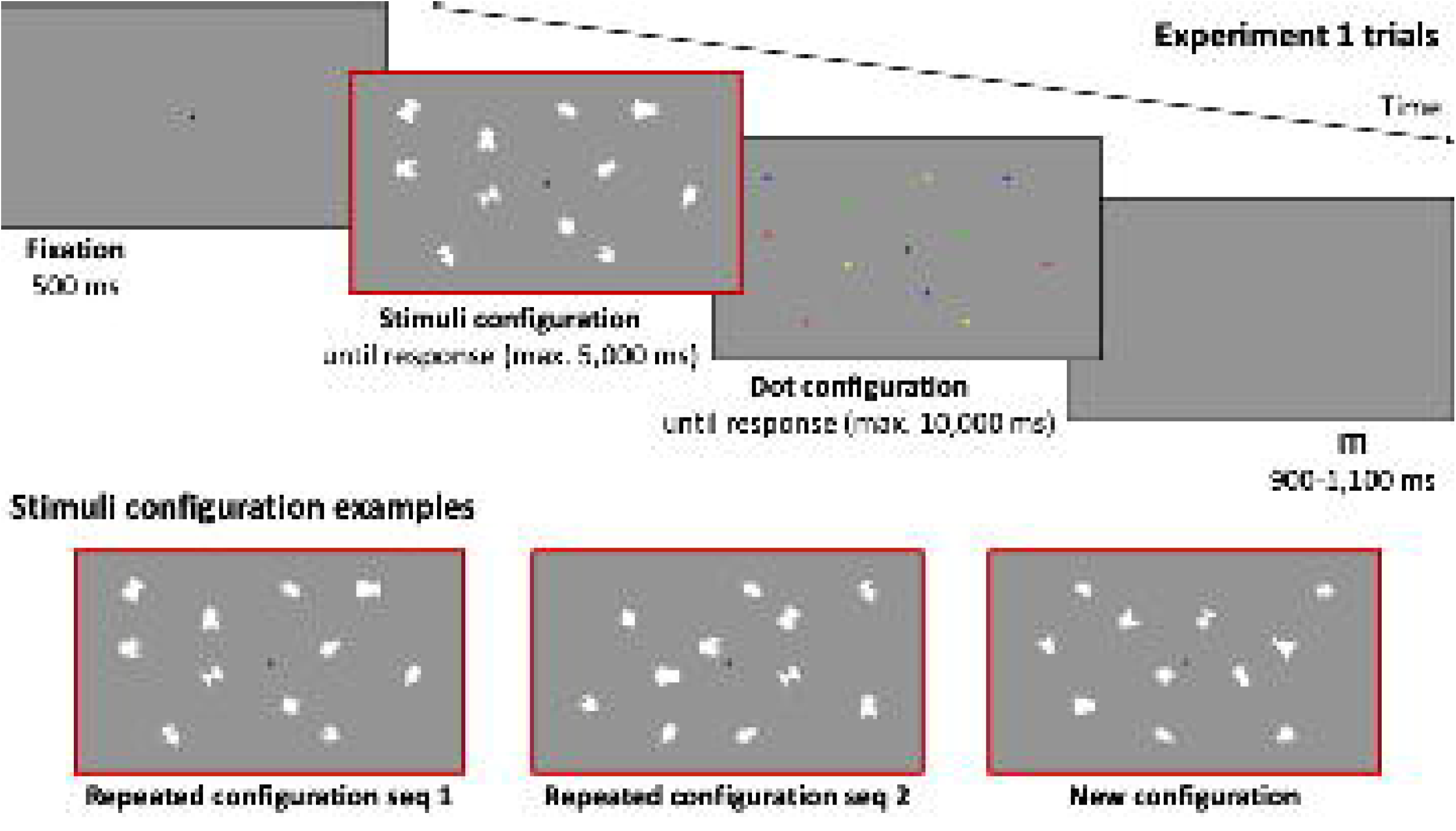
The experimental design of the object cueing paradigm. The upper row displays the schema of a trial. The ‘Stimuli configuration’ shows the array where participants were required to find the stimulus that was symmetric along the vertical axis. After pressing the button, a dot configuration appeared, prompting participants to move the cursor and click on the dot corresponding to the location of the target stimulus in the previous stimulus configuration. The lower row presents possible stimulus configurations: the first two configurations contain the same objects (REPEATED configuration), while the third configuration features different objects from those previously shown (NEW configuration).

The experiment started with 1 to 3 practice sequences (16 trials per sequence, using different stimuli than those in the experimental trials, with the number of practices based on performance) followed by 6 experimental blocks, with each block containing 4 sequences. Each sequence comprised 8 REPEATED trials and 8 NEW trials. Every trial started with a black fixation point (.07° x .07°) which was presented for 500 ms, followed by the stimuli array. Participants were instructed to press the left mouse button with their right hand as quickly as possible when they found the single stimulus that was symmetric along the vertical axis, but they were not informed about any display regularities. The stimulus array was presented for 5,000 ms or until response. After the button press, the array disappeared and was replaced by a screen showing 11 coloured dots (.12° x .12°), each corresponding to the location of a stimulus in the previous display. Participants had to move the cursor over the dot that matched the location of the target stimulus and click to confirm its position. This allowed us to ensure that participants accurately identified the target’s location. This screen was presented for 10,000 ms or until response. The intertrial interval (ITI) varied between 900 and 1,100 ms in 50 ms-long steps. At the end of each sequence, participants received feedback on their reaction time and accuracy.

To test whether participants recognized display regularities, an old/new recognition test was conducted at the end of the session, in which the 8 repeated configurations from the experimental task were presented alongside 8 randomly generated new configurations. The presentation details were the same as before, except that no dot configurations were included in the old-new trials. Participants had to determine whether they recognized the presented configuration from the experimental task. If they recognized it, they clicked the left mouse button with the index finger of their right hand; if they did not, they clicked the right mouse button with their right middle finger. After each response, the cursor automatically returned to the centre of the screen.

### 4.2. RESULTS

Mean accuracy was above 94% on average, but the error rate showed a *Block* main effect, F(5,120) = 6.85, p = .001, η^2^_p_ = .22. Post hoc tests revealed more errors in Block 1 compared to Blocks 4-6 and in Block 2 compared to Blocks 5-6 (all ps ≤ .02). There was no significant main effect of *Configuration*, nor was there a *Block x Configuration* interaction for the error rate.

The mean RT data across the 6 Blocks are presented in Figure 2. Although there was a *Block* main effect, F(3.39,81.42) = 31.41, p < .001, η^2^_p_ = .57, with longer RTs in Block 1 compared to Blocks 2-6, in Block 2 compared to Blocks 3-6, and in Block 3 compared to Block 6 (all ps ≤ .02), neither the *Configuration* main effect nor the *Block x Configuration* interaction reached significance.

**Figure 2.**
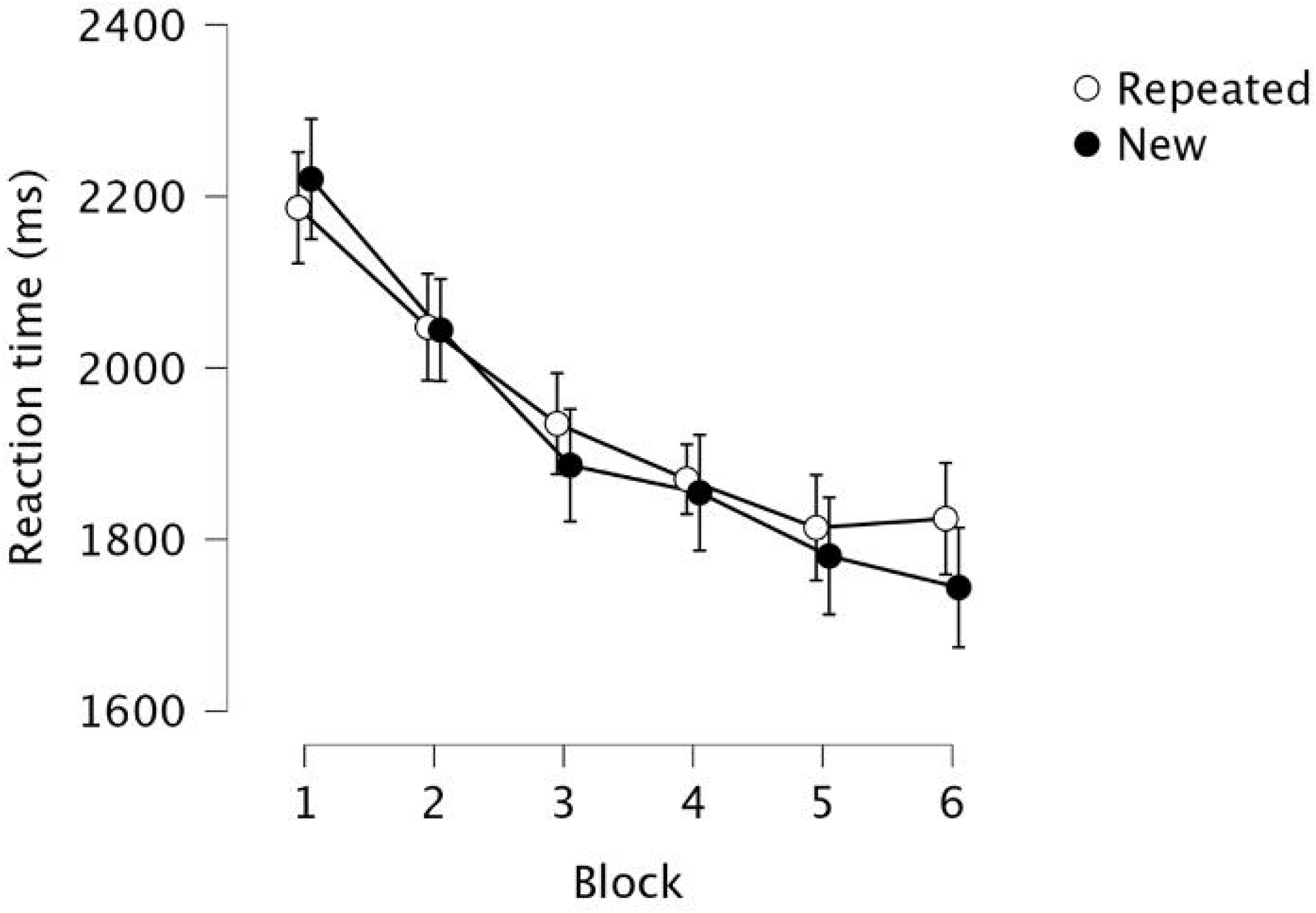
Mean reaction time in milliseconds across the 6 blocks for REPEATED and NEW configurations in Experiment 1. Error bars represent the standard error.

In the old/new recognition test, the discrimination index (Pr) was -.11 (SD = .31), t(24) = -1.80, p = .085, while the bias index (Br) was .52 (SD = .08), t(24) = 1.34, p = .193.

Based on Chun and Jiang (1999), the contextual cueing effect was expected to emerge by Block 2, but when analysing the average RT difference between the NEW and REPEATED configurations at the individual level across Blocks 2-6, we found that only 8 participants exhibited a contextual cueing effect. In contrast, the remaining 17 participants did not show this effect (Figure 3). The analysis in which we included these two groups as a between-subjects factor emerged as a post hoc idea that we explored subsequently.

**Figure 3.**
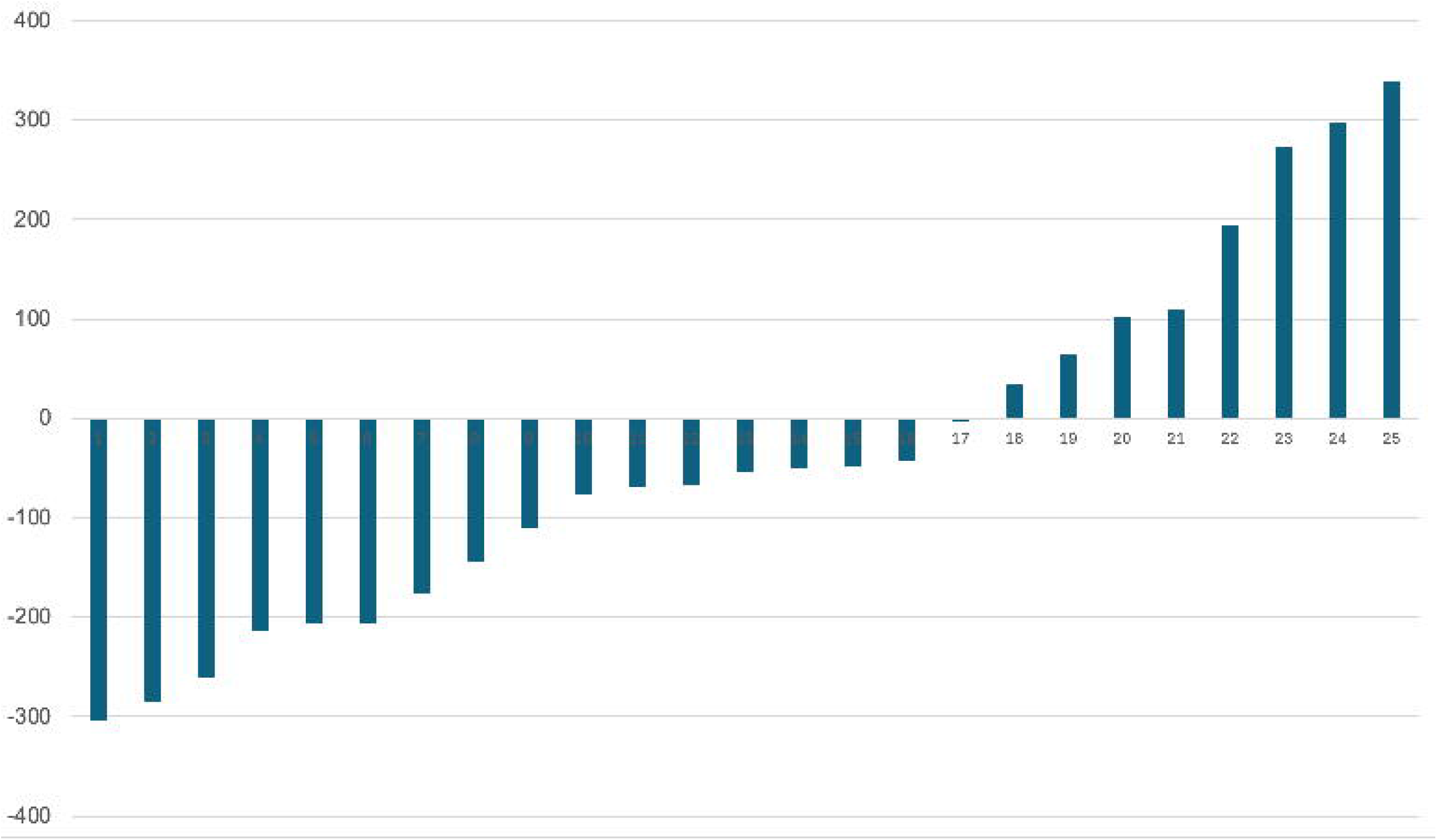
Contextual cueing effect for each participant in Experiment 1, calculated as the average RT difference between NEW and REPEATED configurations across Blocks 2-6. Positive values indicate faster responses to REPEATED configurations.

Based on the main effect of *Configuration* observed in Experiment 1 of Chun and Jiang (1999), we estimated the required sample size to detect a significant effect. Using G*Power with 80% power and an alpha level of 0.05, we determined that a sample size of 8 participants was necessary. Consequently, we could divide our sample into two groups: one where the contextual cueing effect did not emerge (NO CC GROUP) and one where the effect was observed (CC GROUP), and we included *Group* as a between-subjects factor into the Repeated measures of ANOVA.

In Experiment 1, we found that only 8 participants showed a positive contextual cueing effect, that is, the average RT difference in Blocks 2-6 between the NEW and REPEATED conditions exceeded zero. In contrast, the remaining 17 participants did not show this effect. Analysing mean RT, we found a *Configuration x Group* interaction F(1,23) = 47.52, p < .001, η^2^_p_ = .52 (see Figure 4). Post hoc tests revealed that the RT was longer for REPEATED configurations compared to NEW configurations in the NO CC GROUP (p < .001), whereas in the CC GROUP, the opposite pattern was observed: RT was longer for NEW configurations compared to REPEATED configurations (p < .001). The post hoc test of the *Block x Configuration x Group* interaction, F(5,115) = 2.43, p = .039, η^2^_p_ = .10, showed that these differences were present in Blocks 3 and 6 (p = .006 and .020) in the NO CC GROUP and in Block 3 (p = .012) in the CC GROUP.

**Figure 4.**
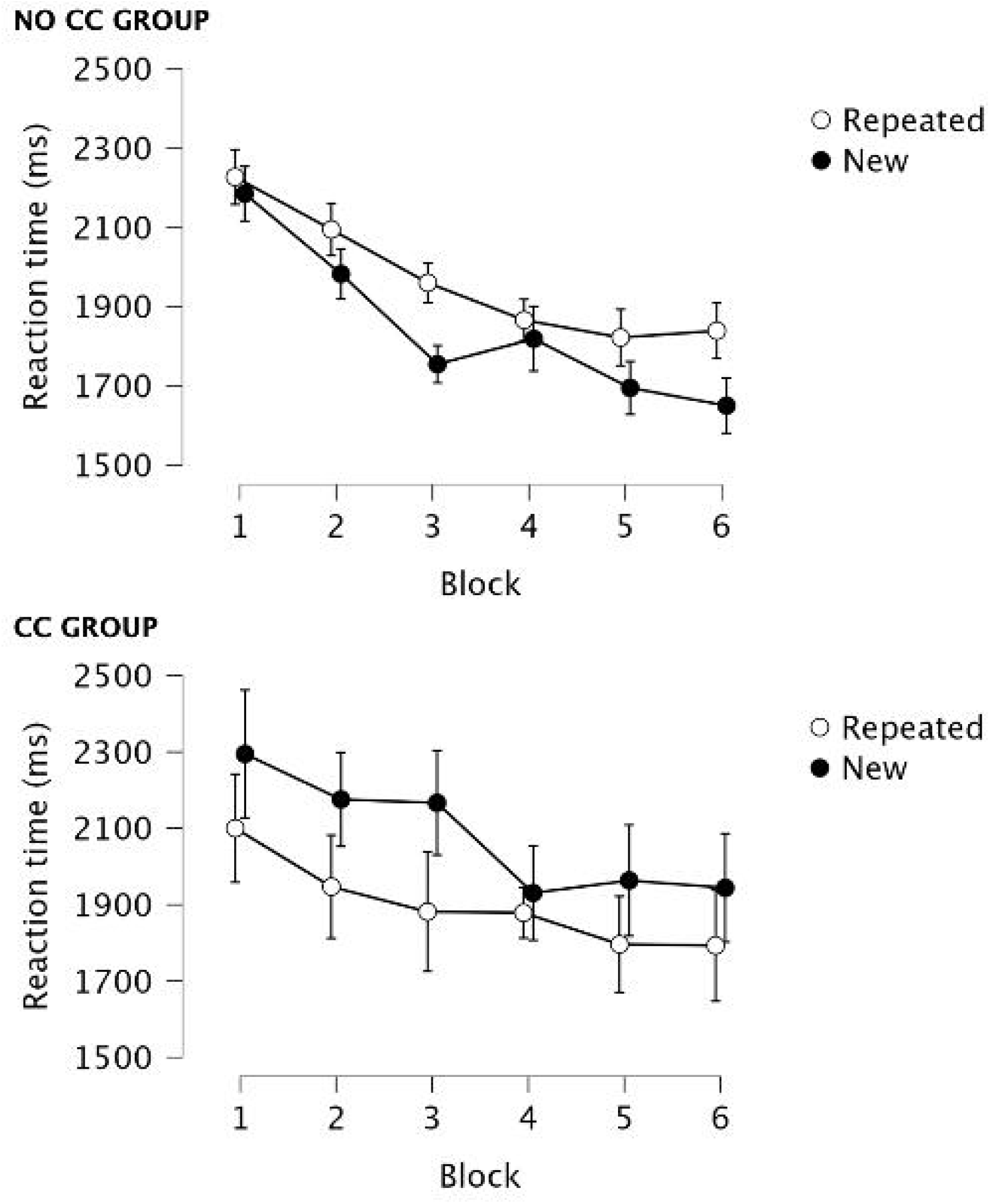
Mean reaction time (RT) in milliseconds across the 6 blocks for REPEATED and NEW configurations in Experiment 1, comparing the group where the positive contextual cueing effect did not emerge (NO CC GROUP) with the group where the positive effect did appear (CC GROUP). Error bars represent the standard error.

In the old/new recognition test, the discrimination index (Pr) was -.16 (SD = .26), t(16) = -2.55, p = .021, and the bias index (Br) was .53 (SD = .07), t(16) = 2.19, p = .044 in the NO CC group. In the CC group, the discrimination index (Pr) was .00 (SD = .38), t(7) = .00, p = .999, and the bias index (Br) was .48 (SD = .09), t(7) = -.49, p = .642.

## 5. EXPERIMENT 2

Similar to the above findings, Lleras and von Mühlenen (2004) failed to observe a contextual cueing effect when they attempted to replicate the spatial contextual cueing experiment by Chun and Jiang (1998). They concluded that participants differed in the strategies they used: some adopted an active, others a passive search strategy. This strategic variability, in turn, may explain the absence of the effect in some individuals. To test this, they manipulated strategy use through explicit instructions, and found that the group instructed to use a passive strategy showed a robust contextual cueing effect. Following their approach, in Experiment 2 we replicated Experiment 1 with one key modification: participants were divided into two groups. One group received instructions promoting an active search strategy, while the other group was encouraged to passively let the target “pop out.”

### 5.1. METHODS

Participants were randomly assigned to two groups. The ACTIVE group included 20 participants (18 female, M = 21.8 years, SD = 2.1, all right-handed), and the PASSIVE group also included 20 participants (15 female, M = 21.7 years, SD = 2.0).

The task was identical to that used in Experiment 1, except that the two groups received different instructions, both verbally and in writing. Based on Lleras and von Mühlenen (2004), the strategy parts of the instructions were as follows (in Hungarian):

Active search strategy (ACTIVE group): “The best strategy for this task, and the one that we want you to use in this study, is to be as active as possible and to ‘search’ for the item as you look at the screen. The idea is to deliberately direct your attention to determine your response. Sometimes people find it difficult or strange to ‘direct their attention’, but we would like you to try your best. Try to respond as quickly and accurately as you can while using this strategy. Remember, it is very critical for this experiment that you actively search for the unique item.”

Passive search strategy (PASSIVE group): “The best strategy for this task, and the one that we want you to use in this study, is to be as receptive as possible and let the unique item ‘pop’ into your mind as you look at the screen. The idea is to let the display and your intuition determine your response. Sometimes people find it difficult or strange to tune into their ‘gut feelings’, but we would like you to try your best. Try to respond as quickly and accurately as you can while using this strategy. Remember, it is very critical for this experiment that you let the unique item just ‘pop’ into your mind.”

### 5.2. RESULTS

Mean accuracy was above 94% on average during the experiment. The error rate in the ACTIVE group was M = 0.061, SD = 0.051, and in the PASSIVE group M = 0.058, SD = 0.044. The error rate decreased as the task progressed, as a *Block* main effect showed, F(3.437, 153.451) = 9.754, p < .001, η^2^_p_ = .64. Post hoc tests revealed more errors in Block 1 compared to Blocks 2-6, and in Block 3 compared to Block 6 (all ps < .035). No other significant effects were detected for the error rate.

The mean RT data for the two groups across the six blocks are presented in Figure 5. A significant main effect of *Block* was observed, F(5,190) = 9.754, p < .001, η^2^_p_ = .20, indicating decreasing RTs over time. Specifically, RTs were longer in Block 1 compared to Blocks 2-6, in Block 2 compared to Blocks 3-6, in Blocks 3 and 4 compared to Blocks 5-6, and in Block 5 compared to Block 6 (all ps < .015). There was no significant main effect of *Configuration*, nor a *Configuration x Block* interaction. Importantly, no main effect of *Group* or interaction involving *Group* was found, indicating that the contextual cueing effect did not differ between groups receiving different instructions. However, consistent with the findings of Experiment 1, some participants exhibited a contextual cueing effect, while others did not. Specifically, for 8 participants in the ACTIVE group and 11 in the PASSIVE group, the average RT difference between NEW and REPEATED configurations across Blocks 2-6 did not exceed 0. In contrast, 12 participants in the ACTIVE group and 9 in the PASSIVE group did show the contextual cueing effect (Figure 6).

**Figure 5.**
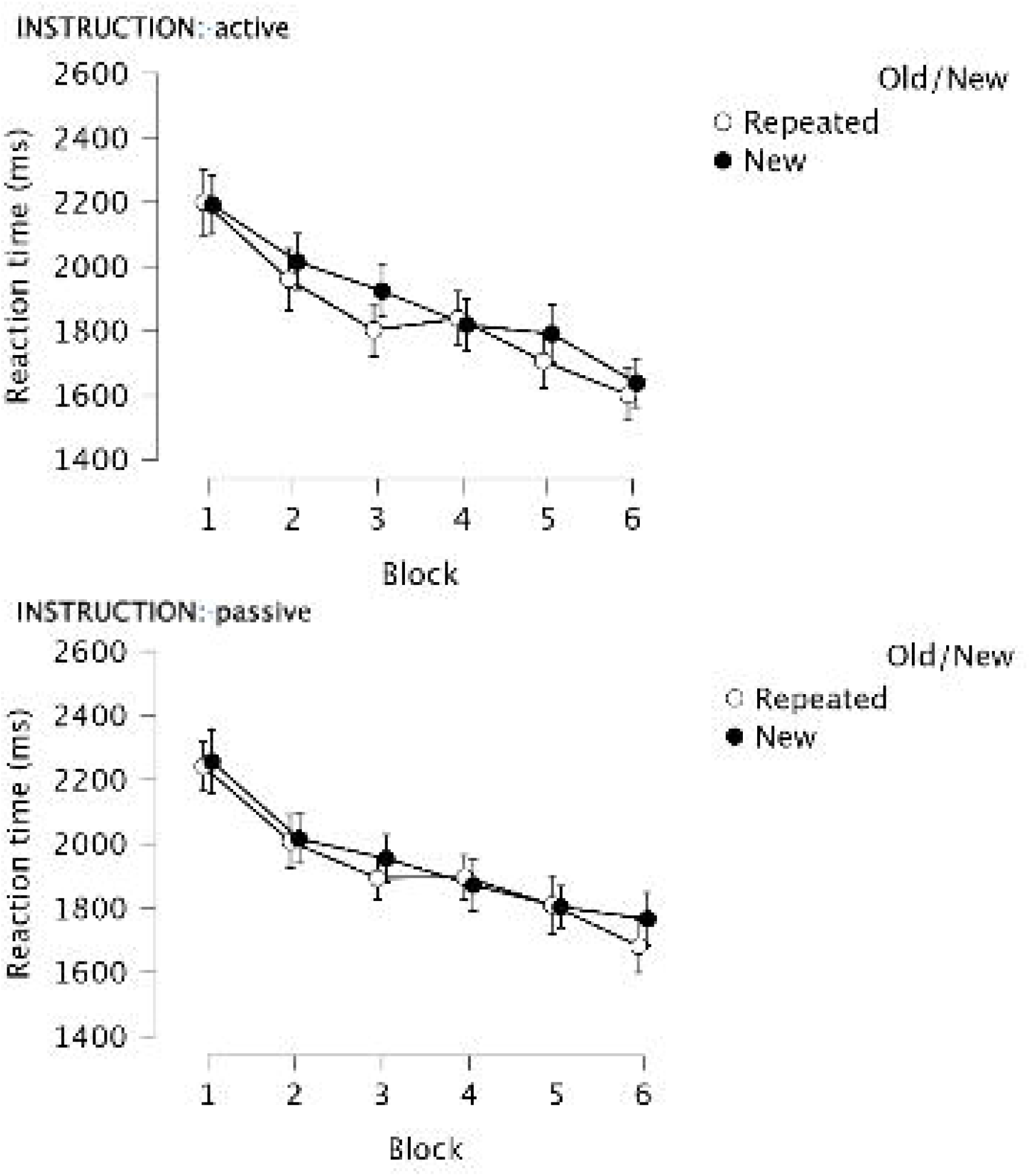
Mean reaction time (RT) in milliseconds across the 6 blocks for REPEATED and NEW configurations in Experiment 2, comparing the two groups receiving different instructions (ACTIVE and PASSIVE). Error bars represent the standard error.

**Figure 6.**
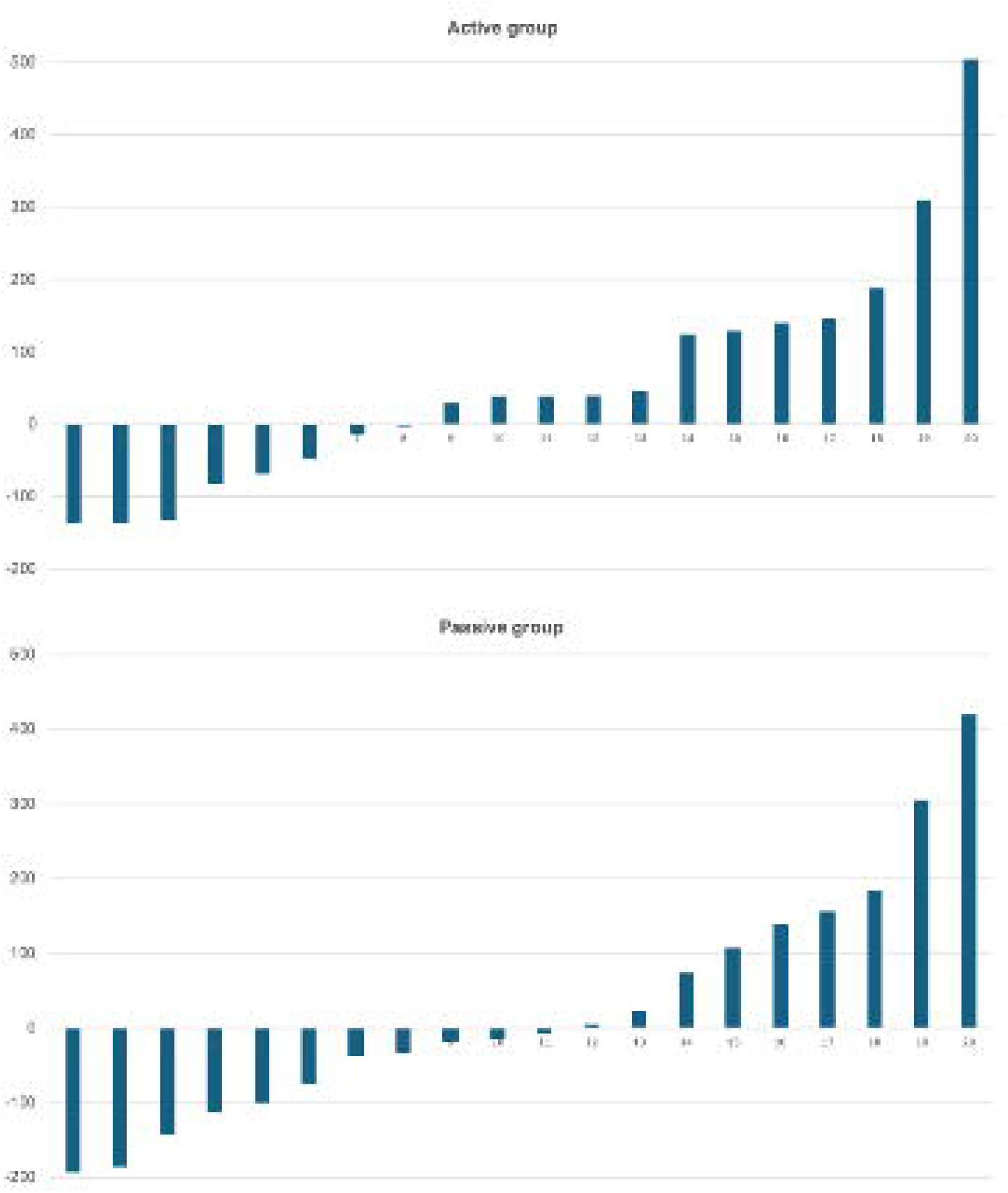
Contextual cueing effect (in ms) in Experiment 2. The upper panel shows results from the ACTIVE group, and the lower panel shows results from the PASSIVE group, separately for each participant, averaged across Blocks 2-6.

In the old/new recognition test, the discrimination index (Pr) was -.05 (SD = .23), t(19) = -.96, p = .348, in the ACTIVE group, and -.01 (SD = .27), t(19) = -.18, p = .857, in the PASSIVE group. The bias index (Br) was .54 (SD = .11), t(19) = 1.72, p = .102, in the ACTIVE group, and .49 (SD = .17), t(19) = -.23, p = .818, in the PASSIVE group. These results are consistent with participants’ subjective reports following the experiment. Although all participants reported that they were able to follow the instructed strategy, none of them in either group noticed that some of the trials were repeated.

## 6. DISCUSSION

Our surroundings are constantly overloaded with stimuli, so it is crucial to extract their regularities in order to support and enhance everyday performance. These stimuli often act as attention distractors, increasing working memory load – particularly challenging for individuals with impaired working memory, such as older adults (Weeks & Hasher, 2018). However, through repeated exposure and learning, they can also aid in controlling behaviour. To better understand this phenomenon, in our study we focused on how object co-occurrences can modify visual search performance.

To investigate how associations form between a target stimulus and surrounding distractors, Chun and Jiang (1999) developed the object contextual cueing paradigm. This paradigm is a variant of the visual search task in which, on some trials, the set of distractors repeats, while on others, a novel set of distractors appears. The paradigm is particularly valuable because it allows researchers to examine from scratch how such associations develop between unfamiliar objects – in this case, symmetrical shapes. Our goal was to replicate this experiment and examine whether repeated contextual configurations facilitate target recognition compared to novel ones as trials progress.

Our results showed that participants responded faster and made fewer errors over time. However, we found no significant difference between the repeated and new configurations. Thus, we were unable to replicate Chun and Jiang’s (1999) findings on object contextual cueing. The inefficiency of repeated object identity information for guiding visual attention and enhancing visual search performance was reported by a previous study as well using everyday objects (Makovski, 2016).

When studying implicit learning, a key question is whether participants are aware of the repeated stimuli. To assess this, we conducted an old/new recognition test. The discrimination and bias indices in both Experiment 1 and 2 indicated that participants did not recognize the repeated configurations, as their decisions were made at chance level. It should be noted, however, that our interpretation of the old/new recognition test is limited due to the small number of trials included (8 previously presented repeated configurations and 8 newly generated ones), which may not provide sufficient statistical power (Smyth & Shanks, 2008).

In Experiment 1, we closely followed the original study except for the response method. While Chun and Jiang replaced the symmetrical shapes with letters, requiring participants to type the letter that appeared in the target’s location, we replaced the shapes with coloured dots, and participants had to click on the dot at the target’s location using a mouse. However, it is unlikely that this modification was responsible for the absence of the effect. One potential concern could be the sample size, but our experiments included 25 and 2×20 participants, respectively, compared to only 17 in the original study. Moreover, based on the results reported by Chun and Jiang, a G*Power analysis with 80% power and an alpha level of 0.05 indicated that a sample size of just 8 participants would have been sufficient to detect the effect.

When investigating the underlying reasons, it becomes apparent that there are large individual differences: the effect emerges in some individuals but not in others. While the majority did not exhibit a contextual cueing effect – in fact, a substantial negative effect was observed in many participants – a smaller subgroup in both experiments showed faster reaction times in repeated configurations. Notably, previous research on spatial contextual cueing has reported that approximately 30% of participants do not exhibit the effect (Lleras & Von Mühlenen, 2004), which may be attributed to individual differences in implicit learning (Nakamura, 2014).

One possible explanation is that some individuals rely more on global perception, while others adopt a more local perceptual strategy. Unlike spatial cueing – where it has been demonstrated that repeating a smaller area around the target is sufficient to produce the effect, similar to presenting the entire display (Brady & Chun, 2007) – object cueing may require perceiving the full scene to recognize co-occurrences.

Another possibility – one we considered the most likely based on the discrimination and bias index results – was that participants differ in their use of active versus passive strategies. Some individuals may follow a passive strategy, completing the task with minimal conscious effort and acquiring associations implicitly. Others, however, may actively orient their attention for searching patterns, trying to recognize them explicitly. This active approach, with its excessive focus and increased working memory load, may actually interfere with implicit learning. This was supported by the study of Lleras and Von Mühlenen (2004) who detected robust contextual cueing effect with emphasizing passive search strategy in the task instruction but not with emphasizing active search strategy.

To further examine these questions, we conducted an additional experiment in which we manipulated participants’ strategy through instruction. Although we used the same manipulation as Lleras and von Mühlenen (2004), and participants reported that they were able to follow the instructed strategy – either finding the target passively or actively – our manipulation did not lead differences between the two groups. In fact, once again, we observed the same heterogeneous pattern within both groups: some participants showed a positive contextual cueing effect, others showed no effect, and yet others showed a negative effect.

In summary, we were not able to replicate Chun and Jiang’s (1999) object contextual cueing effect. However, this does not call into question the existence of the phenomenon itself. Further research is needed to clarify the factors that determine whether or not the effect emerges in a given individual. It is possible that different underlying mechanisms contribute to this variability. For instance, while the literature has long debated whether spatial contextual cueing in young adults reflects early attentional or later response-related processes, Kojouharova et al. (2023) found that in young adults, early and intermediate processes – such as attention allocation and stimulus categorization – drive the effect, whereas in older adults, it is primarily supported by late response selection mechanisms. Incorporating electrophysiological measures in future studies may help determine whether similar differences are present in object contextual cueing and whether distinct neural mechanisms underlie the effect in individuals who do or do not show it.

The data and materials for both experiments are available at *Anonymised url*.

## ACKNOWLEDGEMENTS

We thank Márta Zimmer for her help. This research was supported by the Ministry of Innovation and Technology of Hungary from the National Research, Development and Innovation Fund (OTKA K 132880).

## Notes

https://gin.g-node.org/gaalzs/FOC.git

### Competing Interest Statement

The authors have declared no competing interest.

## REFERENCES

Brady, T. F., & Chun, M. M. (2007). Spatial constraints on learning in visual search: Modeling contextual cuing. Journal of Experimental Psychology: Human Perception and Performance, 33(4), 798–815. 10.1037/0096-1523.33.4.798

Chun, M. M., & Jiang, Y. (2003). Implicit, long-term spatial contextual memory. Journal of Experimental Psychology: Learning, Memory, and Cognition, 29(2), 224–234. 10.1037/0278-7393.29.2.224

Chun, M.M. & Jiang, Y. (1999). Top-down attentional guidance based on implicit learning of visual covariation. Psychological Science, 10, 360–365. DOI: 10.1111/1467-9280.00168

Chun, M.M. & Jiang, Y. (1998). Contextual cueing: implicit learning and memory of visual context guides spatial attention. Cognitive Psychology, 36, 28–71. 10.1006/cogp.1998.0681

Dal Ben, R. (2023). SHINE_color: Controlling low-level properties of colorful images. MethodsX, 11, 102377. 10.1016/j.mex.2023.102377

Danion, J., Meulemans, T., Kauffmann-Muller, F., Vermaat, H. (2001). Intact implicit learning in schizophrenia. American Journal of Psychiatry, 158(6). 10.1176/appi.ajp.158.6.94

Eldridge, L. L., Masterman, D., & Knowlton, B. J. (2002). Intact implicit habit learning in Alzheimer’s disease. Behavioral Neuroscience, 116(4), 722–726. 10.1037/0735-7044.116.4.722

Goujon, A., Didierjean, A., Thorpe, S. (2015). Investigating implicit statistical learning mechanisms through contextual cueing. Trends in Cognitive Sciences, 19(9), 524–533. DOI: 10.1016/j.tics.2015.07.009

JASP Team. (2025). JASP (Version 0.19.3) [Computer software]. https://jasp-stats.org/

Kojouharova, P., Nagy, B., Czigler, I., & Gaál, Z. A. (2023). Mechanisms of spatial contextual cueing in younger and older adults. Psychophysiology, 60, e14361. 10.1111/psyp.14361

Lleras, A., Von Mühlenen, A. (2004) Spatial context and top-down strategies in visual search. Spatial Vision, 17(4-5), 465–482. DOI: 10.1163/1568568041920113

Makovski, T. (2016). What is the context of contextual cueing? Psychonomic Bulletin & Review, 23, 1982–1988. 10.3758/s13423-016-1058-x

Minear, M., & Park, D. C. (2004). A lifespan database of adult facial stimuli. Behavior Research Methods, Instruments & Computers, 36(4), 630–633. 10.3758/BF03206543

Nagy, B., Kojouharova, P., Protzner, A.B., Gaál, Zs.A. (2024). Investigating the Effect of Contextual Cueing with Face Stimuli on Electrophysiological Measures in Younger and Older Adults. Journal of Cognitive Neuroscience, 36(5), 776–799. 10.1162/jocn_a_02135

Nakamura, D. (2015). Individual Differences in Implicit Learning: Current Problems and Issues for Research. In Z. Jin (Ed.), Exploring Implicit Cognition: Learning, Memory, and Social Cognitive Processes (pp. 61–85). IGI Global Scientific Publishing. 10.4018/978-1-4666-6599-6.ch003

Rathus, J. H., Reber, A. S., Manza, L., & Kushner, M. (1994). Implicit and explicit learning: Differential effects of affective states. Perceptual and Motor Skills, 79(1, Pt 1), 163–184. 10.2466/pms.1994.79.1.163

Reber, P.J. (2013). The neural basis of implicit learning and memory: A review of neuropsychological and neuroimaging research. Neuropsychologia, 51 (10), 2026–2042. 10.1016/j.neuropsychologia.2013.06.019

Robin, J., & Moscovitch, M. (2017). Familiar real-world spatial cues provide memory benefits in older and younger adults. Psychology and Aging, 32(3), 210–219. 10.1037/pag0000162

Seger, C. A. (1994). Implicit learning. Psychological Bulletin, 115(2), 163–196. 10.1037/0033-2909.115.2.163

Smyth, A. C., & Shanks, D. R. (2008). Awareness in contextual cuing with extended and concurrent explicit tests. Memory & Cognition, 36, 403–415. 10.3758/mc.36.2.403

Vickery, T. J., Sussman, R. S., & Jiang, Y. V. (2010). Spatial context learning survives interference from working memory load. Journal of Experimental Psychology: Human Perception and Performance, 36(6), 1358–1371. 10.1037/a0020558

Zellin, M., von Mühlenen, A., Müller, H. J., & Conci, M. (2014). Long-term adaptation to change in implicit contextual learning. Psychonomic Bulletin & Review, 21(4), 1073–1079. 10.3758/s13423-013-0568-z

